# The genome of a vestimentiferan tubeworm (*Ridgeia piscesae*) provides insights into its adaptation to a deep-sea environment

**DOI:** 10.1101/2022.08.16.504205

**Authors:** Muhua Wang, Lingwei Ruan, Meng Liu, Zixuan Liu, Jian He, Long Zhang, Yuanyuan Wang, Hong Shi, Mingliang Chen, Feng Yang, Runying Zeng, Jianguo He, Changjun Guo, Jianming Chen

**Affiliations:** State Key Laboratory for Biocontrol, Southern Marine Science and Engineering Guangdong Laboratory (Zhuhai), China-ASEAN Belt and Road Joint Laboratory on Mariculture Technology, School of Marine Sciences, Sun Yat-sen University, Zhuhai 519000, China; State Key Laboratory Breeding Base of Marine Genetic Resources, Key Laboratory of Marine Genetic Resources of Ministry of Natural Resources, Third Institute of Oceanography, Ministry of Natural Resources, Fujian Key Laboratory of Marine Genetic Resources, Xiamen 361005, China; Novogene Bioinformatics Institute, Beijing 100083, China; Guangdong Province Key Laboratory for Aquatic Economic Animals, School of Life Sciences, Sun Yat-sen University, Guangzhou 510275, China; Maoming Branch, Guangdong Laboratory for Lingnan Modern Agricultural Science and Technology, Maoming 525435, China; Fujian Key Laboratory on Conservation and Sustainable Utilization of Marine Biodiversity, Fuzhou Institute of Oceanography, Minjiang University, Fuzhou, 350108, China

**Author notes:** These authors contributed equally to this work (Muhua Wang, Lingwei Ruan, Meng Liu). **Correspondence:** Jianming Chen, Changjun Guo, Jianguo He.

**Keywords:** vestimentiferan tubeworm, *Ridgeia piscesae*, genome, deep-sea adaptation, lifespan

## Abstract

Vestimentifera (Polychaeta, Siboglinidae) is a taxon of deep-sea worm-like animals living in the deep-sea hydrothermal vent and cold seep areas. The morphology and lifespan of *Ridgeia piscesae,* which is the only vestimentiferan tubeworm species found in the hydrothermal vents on the Juan de Fuca Ridge, vary greatly according to the endemic environments. Recent analyses have revealed the genomic basis of adaptation in three vent- and seep-dwelling vestimentiferan tubeworms. However, the evolutionary history and mechanism of adaptation in *R. piscesae*, a unique species in the family *Siboglinidae*, is remained to be investigated. Here we report a high-quality genome of *R. piscesae* collected at Cathedral vent of the Juan de Fuca Ridge. Comparative genomic analysis revealed that that the high growth rates of vent-dwelling tubeworms might derive from small genome size. The small genome sizes of these tubeworms are attributed to the repeat content but not the number of genes and intron sizes. Additionally, four genes involved in cell proliferation were subject to positive selection in the genome of *R. piscesae*, suggesting that, besides apoptosis, cell proliferation is important for regulating growth rate in this species.

## Introduction

The discovery of deep-sea hydrothermal vents and cold seeps, as well as their associated ecosystems have revolutionized our view of biology and understanding of the energy sources that fuel primary productivity on Earth (Corliss et al., 1979; Paull et al., 1984; Petersen et al., 2011). Hydrothermal vents are areas on the ocean floor where hot, anoxic, chemical-rich water is expelled into the cold, oxygen-rich deep ocean (Von Damm, 1995). Cold seeps are areas where methane, hydrogen sulfide, and other hydrocarbons seep or emanate as gas from deep geologic sources (Suess, 2014). Both hydrothermal vents and cold seeps are characterized by high hydrostatic pressure, darkness, lack of oxygen and photosynthesis-derived nutrients, and high concentrations of toxic chemicals (Levin, 2005). Organisms inhabiting around hydrothermal vents and cold seeps develop unique characters to adapt to the adverse conditions (Grassle, 1985; Childress and Fisher, 1992). Due to the complete absence of light, hydrothermal vent and cold seep ecosystems are driven by the chemosynthesis instead of photosynthesis (Stewart et al., 2005; Vanreusel et al., 2009). The process is completed by the chemosynthetic microorganisms, which cooperate with a variety of macrobenthos to form chemosynthetic symbioses and contribute to the primary production supporting the ecosystem (Dick, 2019).

Vestimentifera (Polychaeta, Siboglinidae) is a taxon of deep-sea worm-like animals living in the deep-sea hydrothermal vent and cold seep areas (Bright and Lallier, 2010). The body of adult vestimentiferan tubeworm is enclosed in a chitinous tube that is closed at the posterior end. Vestimentiferan tubeworms lack a digestive tract and rely on symbiosis with chemoautotrophic microorganisms, which inhabit in an specialized internal organ, to derive their metabolic needs (Vrijenhoek, 2010). The first discovery of chemoautotrophic symbiont in *Riftia pachyptila,* a vestimentiferan tubeworm inhabiting around hydrothermal vents on the East Pacific Rise (EPR), initiated the intensive study of these deep-sea tubeworms (Corliss et al., 1979). The body of adult tubeworm is comprised of four main parts. The anteriorly located branchial plume is the primary site of gas exchange with the environment. Below the plume is the vestimentum, where heart, gonopores and a simplified brain are located. The trophosome, which is primarily composed of symbiont-containing bacteriocytes and blood vessels, is located below vestimentum. And the segmented opisthosoma is located below vestimentum (Jones, 1981; Hand, 1987; Schulze, 2001).

Lifespan varies greatly between the vent- and seep-dwelling tubeworms (Lutz and Kennish, 1993). The lifespan of *R. pachyptila,* which thrives in relatively strong and continuous diffuse hydrothermal flow, was estimated to be less than 10 years (Urcuyo et al., 2007). In contrast, *Lamellibrachia luymesi,* which live around cold seeps in the Gulf of Mexico, can live up to 250 years (Bergquist et al., 2000). In general, vent-dwelling tubeworms grow faster than seep-dwelling tubeworms. The growth rate of *R. pachyptila* can reach up to about 160 cm yr^-1^, while the growth rate of *L. luymesi* is only about 3 cm yr^-1^ (Shank et al., 1998; Thiebaut et al., 2002; Fisher et al., 2008). Previous analysis revealed that both vent- and seep-dwelling tubeworms have high rates of cell proliferation. Recent analysis suggested that the variation of growth rates is attributed to the variation of apoptosis between vent- and seep-dwelling tubeworms, where apoptosis is substantially downregulated in vent-dwelling species (Pflugfelder et al., 2009).

The Juan de Fuca Ridge in the northeast Pacific Ocean is characterized by a broad heterogeneity of chemical environments, ranging from vigorous, high-temperature vents to diffuse flow (Tivey et al., 1999). The hydrothermal vents on the Juan de Fuca Ridge provide numerous biotic habitats, which support the growth of a large quantity of endemic organisms (Tsurumi and Tunnicliffe, 2003). Although the biomass of endemic hydrothermal vent fauna is high, there is only a few macrofauna species dominating a particular vent community (Lutz et al., 1998; Tunnicliffe et al., 1998). *Ridgeia piscesae* Jones (1985) is the only vestimentiferan tubeworm species in the hydrothermal vents on the Juan de Fuca Ridge. The tubeworm occurs at high density in most of the vents, acts as an ecosystem-structuring species by providing habitats for several other organisms and serving as a primary producer through chemosynthetic endosymbiosis (Childress and Fisher, 1992; Urcuyo et al., 2003).

*Ridgeia piscesae* adopts different strategies to adapt to diverse environments in the vents on Juan de Fuca Ridge. First, a strong phenotypic plasticity, where a given genotype expresses different phenotypes in different ecological setting, was identified in *R. piscesae* (Carney et al., 2007). Two extreme growth forms (morphotypes) of *R. piscesae,* “short-fat” and “long-skinny”, were discovered in the geologically and chemically diverse vent fields (Southward et al., 1995). “Short-fat” *R. piscesae* prefers relatively high flow vent fluid of high temperature (up to 30°C) and high concentrations of sulfide, while “long-skinny” morphotype adapts to ambient temperature (2°C) and low concentrations of sulfide at areas of diffuse hydrothermal fluids (Carney et al., 2007). These two morphotypes diverge in several morphological characters (Jones, 1987). The tube of “short-fat” morphotype has a generally constant diameter of 2-3 centimeters, while the tube diameter of “long-skinny” morphotype reduces from the anterior to posterior. This morphotype acquires sulfide by their buried posterior tube sections in areas where sulfide is generally not detectable around their plumes (Urcuyo et al., 2003). Thus, unlike other vent-dwelling vestimentiferan species, *R. piscesae* can thrive in the areas of diffuse vent fluids. Second, lifespan of *R. piscesae* vary greatly according to the endemic environments (Urcuyo et al., 2007). The species can grow with high growth rates ranging from 6 to 95 cm yr^-1^ under favorable condition, and can grow very slowly when exposed to low levels of vent flow and sulfide (Tunnicliffe et al., 1997; Sarrazin et al., 1998). Strong phenotypic plasticity and flexible lifespan allows *R. piscesae* to adapt to diverse habitats and makes it a unique species in the family *Siboglinidae.* Recent genomic analyses have revealed the genetic basis of adaptation in three vent- and seep-dwelling vestimentiferan tubeworms (Li et al., 2019; Sun et al., 2021; de Oliveira et al., 2022). Although its critical role in supporting the vent ecosystem, the evolutionary history and mechanism of adaptation in *R. piscesae*, a unique deep-sea tubeworm, is remained to be investigated.

Here, we assembled and annotated a high-quality genome of *R. piscesae* collected at Cathedral vent of the Juan de Fuca Ridge. Evolutionary analysis inferred that two vent-dwelling species (*R. piscesae* and *R. pachyptila*) started to diverge due to the separation of the Gorda/Juan de Fuca/Explorer (GFE) ridge systems and the East Pacific Rise (EPR). Comparative genomic analysis revealed that that the high growth rates of vent-dwelling tubeworms might derive from small genome size. The small genome sizes of these tubeworms are attributed to the repeat content but not the number of genes and intron sizes. Additionally, four genes involved in cell proliferation were subject to positive selection in the genome of *R. piscesae,* suggesting that, besides apoptosis, cell proliferation is important for regulating growth rate in this species.

## Results

### Genome assembly and annotation

The samples of *R. piscesae* were collected from the deep-sea hydrothermal vent at Cathedral vent, Main Endeavor Field of the Juan de Fuca Ridge (47° 56’ N, 129° 05’ W, 2,181 m depth). Short-insert paired-end (180 bp, 300 bp and 500 bp) and long-insert mate-pair (2 kb, 5 kb, 10 kb and 15 kb) sequencing libraries were constructed and sequenced on the Illumina HiSeq 2000 platform. A total of 247.74 Gb sequencing data was generated (**Supplementary Table 1**). Based on the *k*-mer distribution of 180 bp paired-end Illumina read, the genome size of was estimated to be 694.79 Mb with a heterozygosity of 1.2% (**Supplementary Fig. S1**). The final assembly of *R. piscesae* genome was 574.96 Mb with a contig N50 size of 10.42 kb and a scaffold N50 size of 230.23 kb (**Supplementary Table S2**).

A total of 87.4% sequencing reads can be aligned unambiguously to the assembled *R. piscesae* genome sequence, covering 99.74% of the assembly (**Supplementary Table S3**). In addition, 99.63% of Trinity assembled sequences (Unigenes) could be aligned to the assembly (**Supplementary Table S4**). The integrity of genome assembly was further assessed using benchmarking universal single-copy orthologs (BUSCO) tools. The result showed that 897 of 978 (92.6%) single-copy metazoan genes (obd10) were captured in the assembly (**Supplementary Table S5**). These results demonstrated that the *R. piscesae* genome is of high quality, and comparable to the previously published tubeworm genomes (**Table 1**) (Li et al., 2019; Sun et al., 2021; de Oliveira et al., 2022).

**Table 1.**
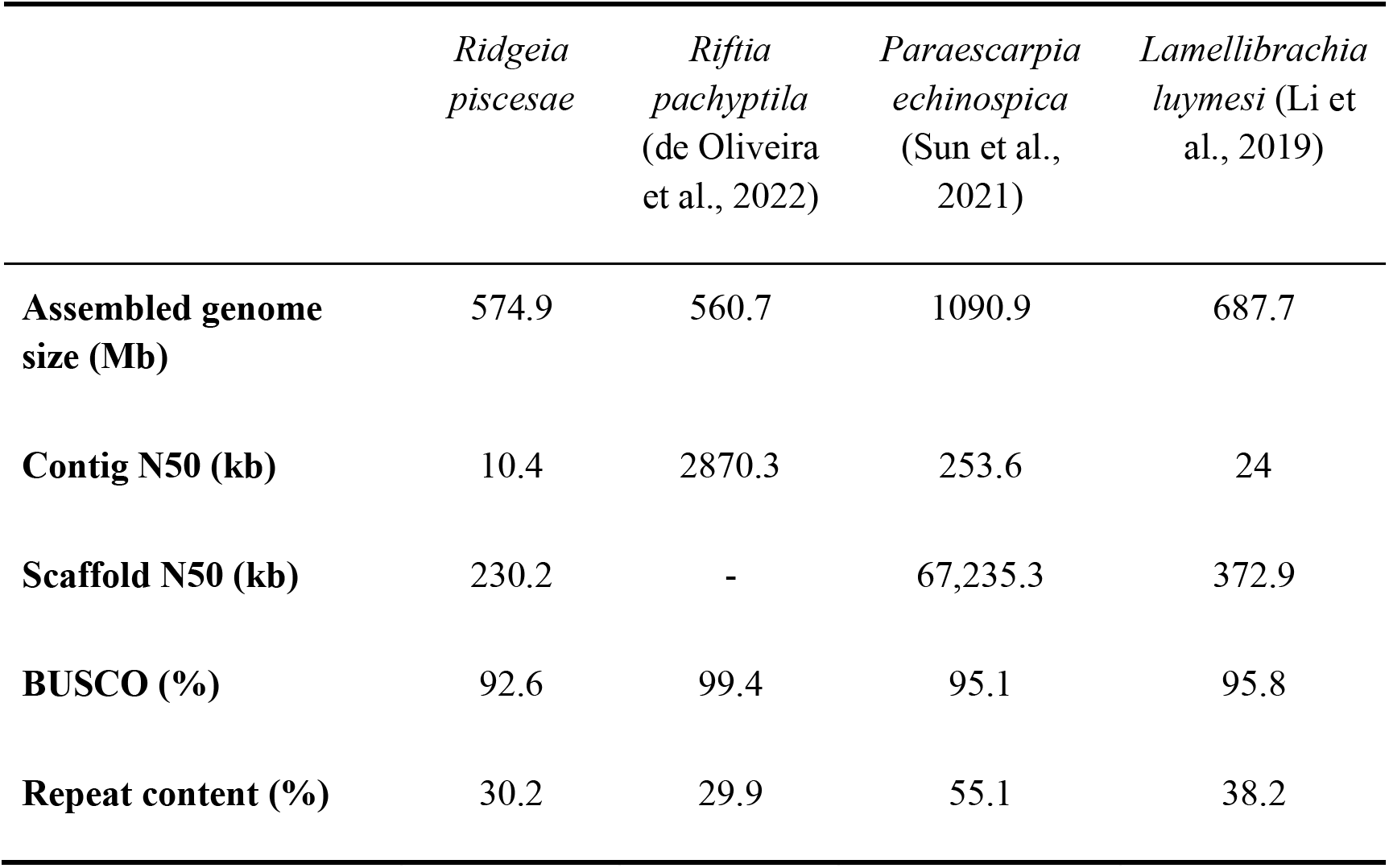
Genome assembly statistics of four deep-sea vestimentiferan tubeworms

|  | *Ridgeia*<br>*piscesae* | *Riftia*<br>*pachyptila*<br>(de Oliveira<br>et al., 2022) | *Paraescarpia*<br>*echinospica*<br>(Sun et al.,<br>2021) | *Lamellibrachia*<br>*luymesii* (Li et<br>al., 2019) |
| --- | --- | --- | --- | --- |
| **Assembled genome**<br>**size (Mb)** | 574.9 | 560.7 | 1090.9 | 687.7 |
| **Contig N50 (kb)** | 10.4 | 2870.3 | 253.6 | 24 |
| **Scaffold N50 (kb)** | 230.2 | - | 67,235.3 | 372.9 |
| **BUSCO (%)** | 92.6 | 99.4 | 95.1 | 95.8 |
| **Repeat content (%)** | 30.2 | 29.9 | 55.1 | 38.2 |

Transposable elements (TEs) accounted for 30.17% in the *R. piscesae* genome assembly, with long interspersed elements (LINEs, 8.08%) as the most abundant class of TEs (**Supplementary Table S6**). The *R. piscesae* genome encodes 24,096 protein-coding genes, of which 95.54% are annotated based on known proteins in diverse public protein databases (**Supplementary Table S7**).

### Phylogenomic analyses

To infer the evolutionary history of *R. piscesae*, a maximum-likelihood (ML) phylogenetic tree was constructed using single-copy orthologs of *R. piscesae* and 14 metazoans with *Adineta vaga* as an outgroup (**Fig. 1, Supplementary Fig. S2, Supplementary Table S8**). Two vent-dwelling tubeworms (*R. piscesae* and *R. pachyptila)* formed a clade. *Paraescarpia echinospica* and *L. luymesi* from cold seeps are basal to the vent clade. These results corroborate the view that vent-dwelling tubeworms might derived from their seep-dwelling relatives (Black et al., 1997; Halanych, 2005).

**Figure 1.**
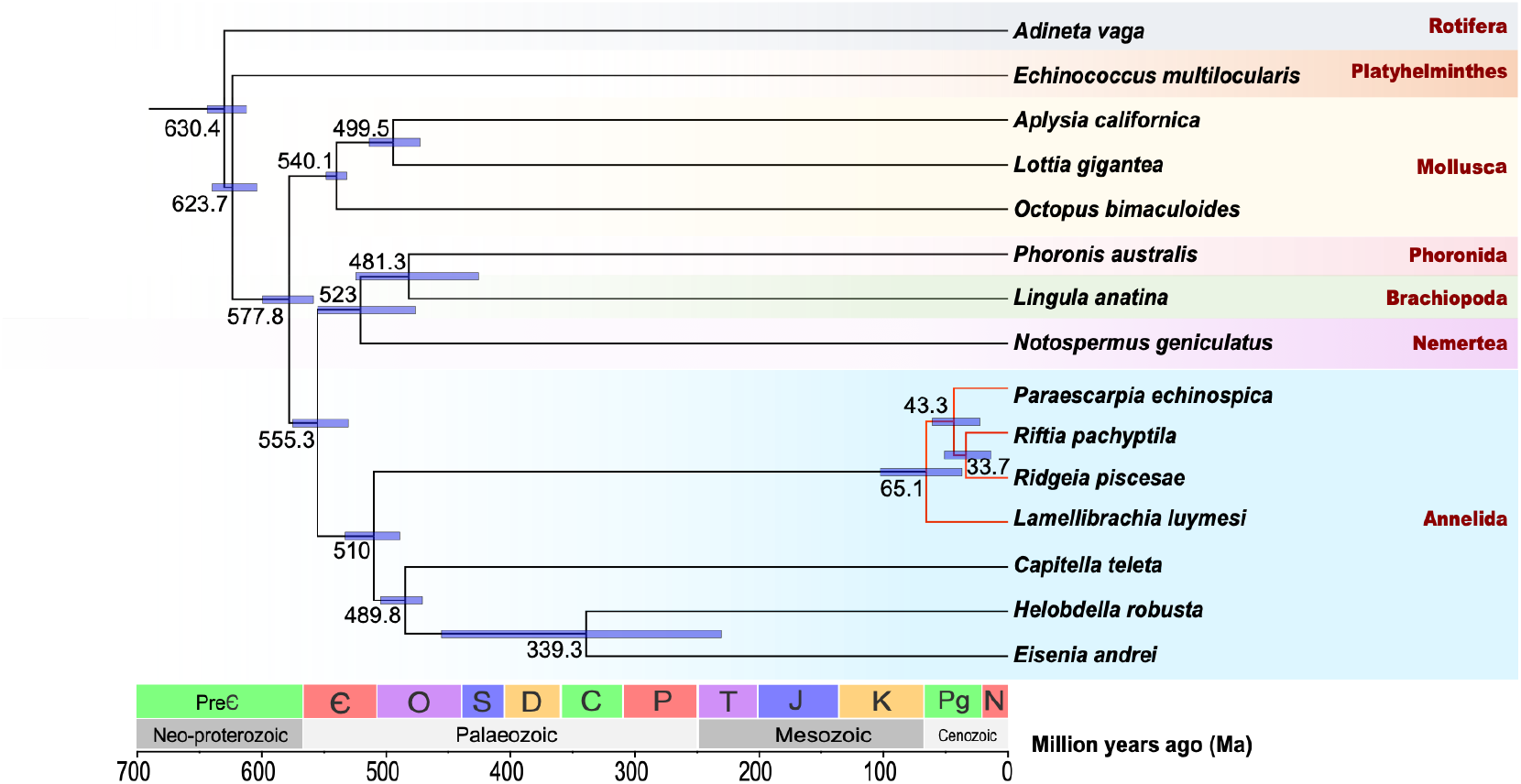
A species tree of *R. piscesae* and 14 metazoans. Single-copy orthologs were used to reconstruct the phylogenetic tree. The divergence time between species pairs was listed above each node, and 95% confidence interval of the estimated divergence time was denoted as blue bar. *R. piscesae* diverged from *R. pachyptila* approximately 33.7 million years ago.

*Ridgeia piscesae* is endemic to the Gorda/Juan de Fuca/Explorer (GFE) ridge systems, and *R. pachyptila* is discovered on the East Pacific Rise (EPR). The subduction of the Farallon-Pacific Ridge separated GFE and EPR between 28.5-35 Ma. Molecular clock analysis revealed that *R. pachyptila* diverged from *R. piscesae* approximately 33.7 million years (Ma). This result suggests these two vent-dwelling species started to diverge due to adaptation to the newly formed ridge systems. The divergence time of *L. luymesi* and other three tubeworms was estimated to be approximately 65.1 Ma, corroborating the view that modern vestimentiferan tubeworms started to diverge during the early Cenozoic Era (Little and Vrijenhoek, 2003; Li et al., 2019; Sun et al., 2021).

#### 3.3 Genome evolution of vestimentiferan tubeworms

It has been demonstrated that several factors, including repeat content, number of genes, and the intron sizes, contributed to the variation of genome sizes among different organisms (Lynch, 2007; Niu et al., 2022). In addition, previous reports proposed that the differences of genome sizes among deep-sea tubeworms might be attributed to the numbers of repetitive elements and genes (Sun et al., 2021; de Oliveira et al., 2022). The assembled size of *R. piscesae* genome is closed to the size of *R. pachyptila,* but smaller than the sizes of two seep-dwelling tubeworms (*L. luymesi* and *P. echinospica)* (**Table 1**). The genomes of cold seep dwelling tubeworms have more transposable elements (TE), especially DNA transposons, LINEs and LTR retrotransposons, than hydrothermal vent dwelling tubeworms (**Fig. 2A**). TEs accounted for 38.2% and 55.1% of *L. luymesi* and *P. echinospica* genomes, and they constituted 30.2% and 29.9% of the genomes of *R. piscesae* and *R. pachyptila.* A strong positive correlation (R^2^ = 0.98, *P* = 0.0052) was identified between genome size and repeat content in these four species (**Fig. 2B**), suggesting that TEs are a major contributor to the genome size evolution in vestimentiferans. Repeat landscape plots indicate that TE activity is different between *R. pachyptila* and three other tubeworm species (**Fig. 3**). There are recent expansions of TEs in the genomes of *L. luymesi, P. echinospica*, and *R. piscesae*, but not in *R. pachyptila*. The main contributor to recent TE expansions in *L. luymesi*, *P. echinospica*, and *R. piscesae* appear to have been LINEs and DNA transposons. Nonetheless, only LINEs were expanded recently in the genome of *R. pachyptila* (**Fig. 3D**).

**Figure 2.**
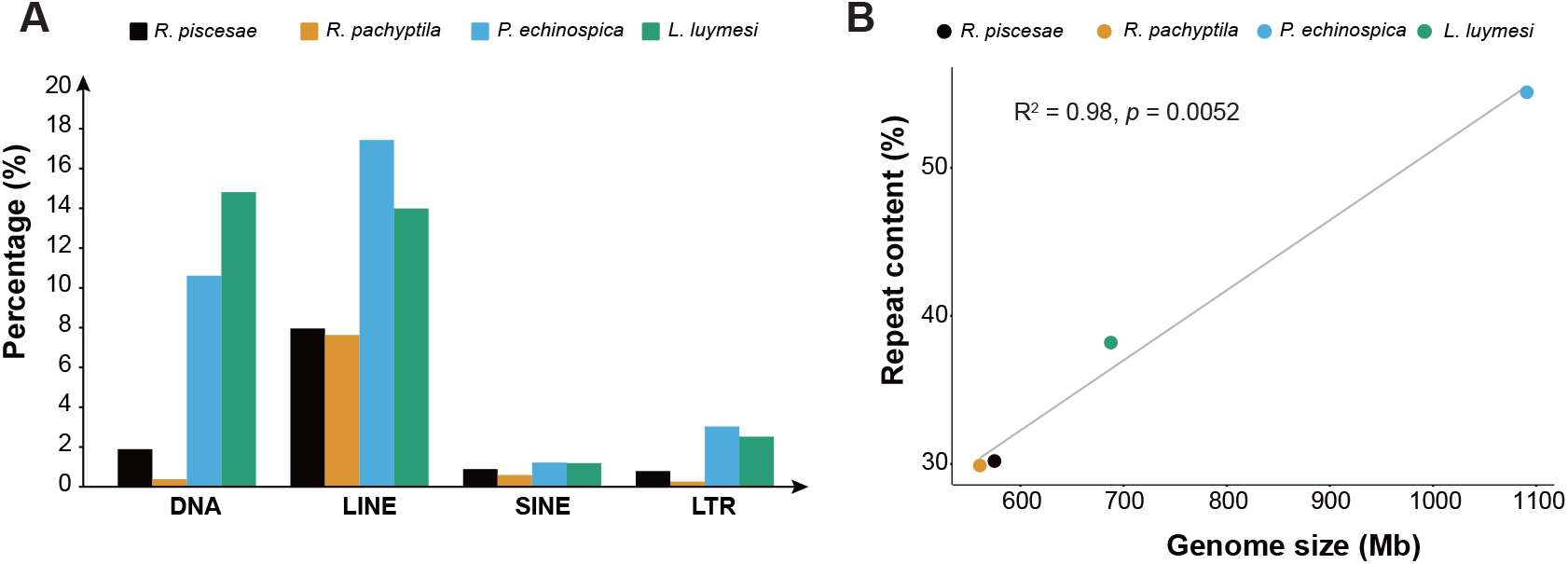
Genome size evolution in four vestimentiferan tubeworms. (**A**) Comparison of the occurrence and composition of repetitive elements in the genomes of 4 vestimentiferan tubeworm. (**B**) The relationship between repeat content and genome size in 4 vestimentiferan tubeworm. A strong positive correlation (R^2^ = 0.98, *P* = 0.0052) was identified between genome size and repeat content in these four species.

**Figure 3.**
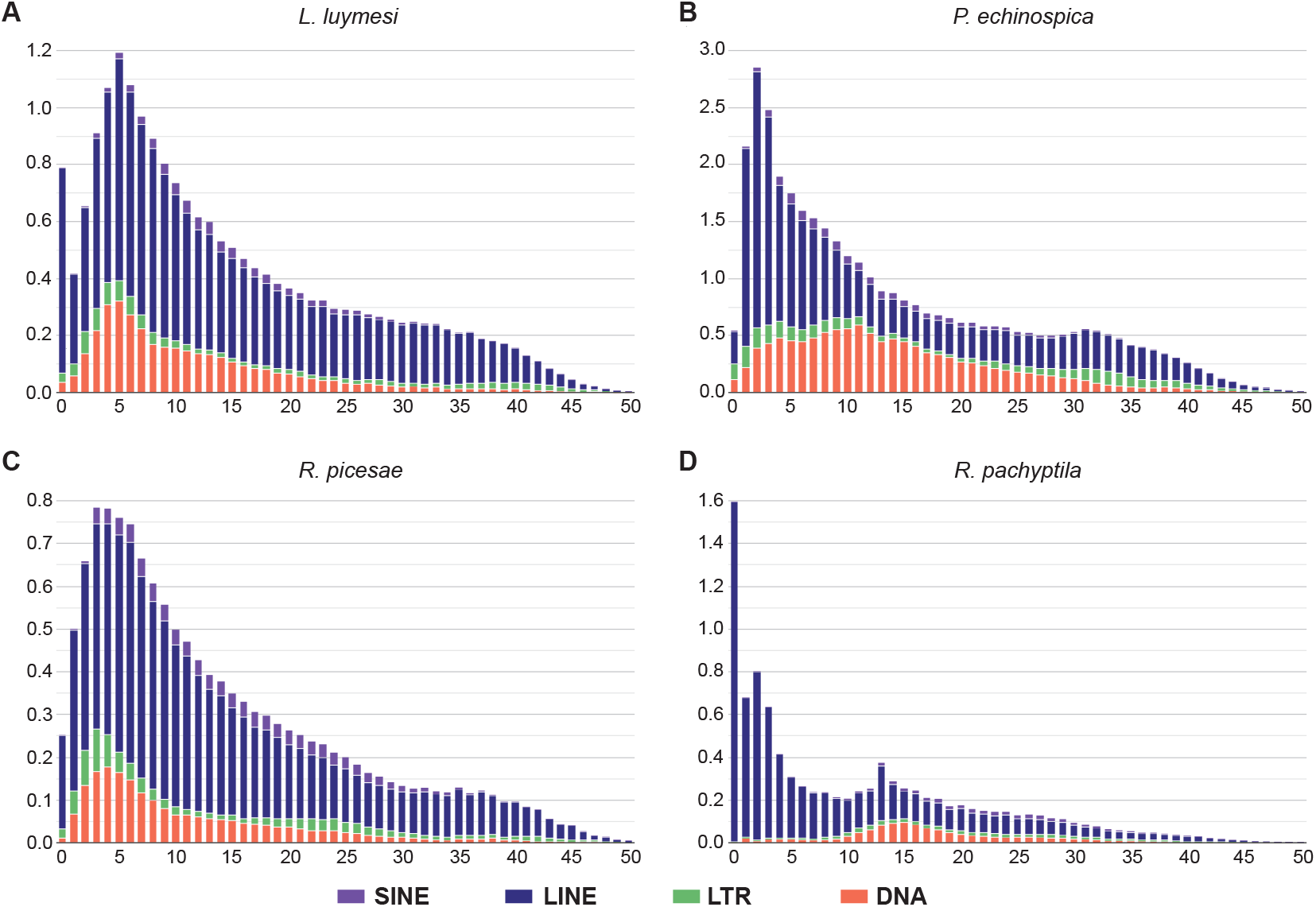
Transposable element-accumulation profile in the genomes of four vestimentiferan tubeworms. There are recent expansions of TEs in the genomes of *L. luymesi*, *P. echinospica*, and *R. piscesae*, but not in *R. pachyptila*.

The number of annotated gene models in *R. piscesae* genome (24,096) is closed to the ones in the genomes of *R. pachyptila* (25,984) and *P. echinospica* (22,642) but less than the ones in *L. luymesi* genome (38,998). Introns account for 220.1 Mb and 204.7 Mb in the genomes of two seep-dwelling tubeworms (*L. luymesi* and *P. echinospica),* as well as 264.8 Mb and 234.5 Mb in the genomes of two vent-dwelling tubeworms (*R. pachtypila* and *R. piscesae)* (**Supplementary Table S9**). Average length of introns in the genome of *R. piscesae* is longer than introns of other three species. Additionally, two vent-dwelling tubeworms with smaller genome sizes have higher ratios of intron / exon length than the seep-dwelling tubeworms. Thus, gene number and intron size do not contribute to the differences of genome sizes between seep- and vent-dwelling tubeworms.

Previous study revealed that *R. pachyptila* experienced reductive evolution with more contracted than expanded gene families in the genome (de Oliveira et al., 2022). Gene-family analysis of four tubeworm species identified a core set of 10,225 gene families (**Fig. 4A**). In total, 601 and 279 lineage-specific gene families were identified in *R. piscesae* and *R. pachyptila*, respectively, which are much less than the ones in *L. luymesi* (1181) and *P. echinospica* (1045). Additionally, gene-family analysis of 12 lophotrochozoans revealed that the numbers of expanded gene families were substantially less than contracted gene families in the two vent-dwelling tubeworms, while more gene families were expanded than contracted in their seep-dwelling counterparts (**Fig. 4B**). These results indicate that the genomes of vent-dwelling tubeworms were characterized by gene loss.

**Figure 4.**
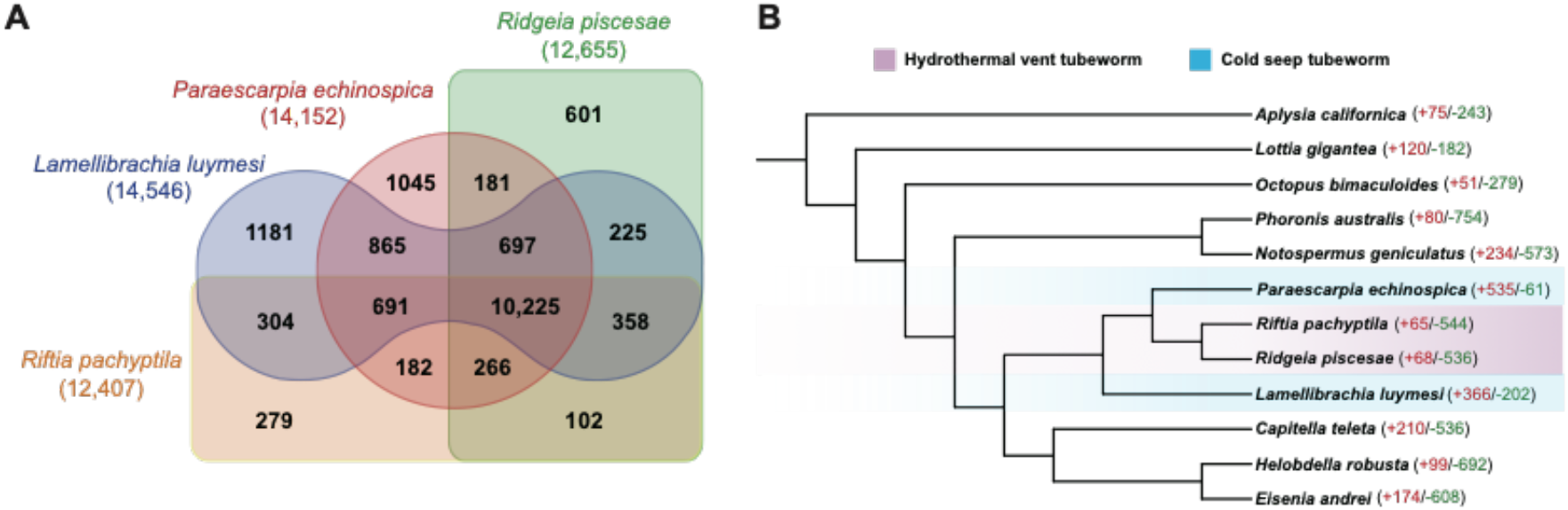
Protein family evolution in four vestimentiferan tubeworms. (**A**) Venn diagram of shared and unique gene families in four vestimentiferan tubeworm species. Lineage-specific gene families of *R. piscesae* and *R. pachyptila* are much less than the ones of *L. luymesi* and *P. echinospica.* (**B**) Gene family expansion/contraction analysis of 4 vestimentiferan tubeworms and 8 other lophotrochozoans.

*Hox* genes are a set of conserved regulators that specify regions of the body plan of an embryo along anterior-posterior axis in metazoans (Pearson et al., 2005). One of the Hox genes (*Antp*) plays a role in the development of posterior segment of several marine annelids (Bakalenko et al., 2013). Loss of *Antp* was apparent across all four tubeworm genomes (**Supplementary Fig. S3**), corroborating the view that the loss of *Antp* contributes to the reduced segmentation of the posterior region of juvenile worms in vestimentiferans (Sun et al., 2021). *Lox2* gene is missing in the genome of *L. luymesi* but presented in the genomes of three other tubeworms, suggesting the loss of this gene might be a lineage-specific event.

#### 3.4 Genomic basis of deep-sea adaptation

Hemoglobins (Hbs) in vestimentiferan tubeworms, which bind oxygen and sulfide simultaneously and provide substrate for chemosynthesis by the symbionts, facilitate the adaptation of these species to deep-sea reducing environments. Four heme-containing chains were identified (A1, A2, B1, B2) in hemoglobins of vestimentiferans (Zal et al., 1997). To elucidate the evolution of Hbs in vestimentiferans, we identified Hb genes in the genomes of four tubeworms species. A single copy of A2 and B2 Hb genes, as well as two copies of A1 genes were identified in each of the tubeworm genomes (**Fig. 5**). Previous studies found that the group of B1 Hbs were significantly expanded in *L. luymesi*, *P. echinospica,* and *R. pachyptila* (Li et al., 2019; Sun et al., 2021; de Oliveira et al., 2022). With 17 identified genes, the group of B1 Hbs was also expanded in the genome of *R. piscesae*. The free cysteine residues in A2 and B2 chains contribute to the sulfide-binding ability of the vestimentiferan Hbs (Bailly et al., 2002). Additionally, the free cysteine was identified at the same position in B1 Hb genes as in A2 Hb genes of *R. piscesae* (**Supplementary Fig. S4**), corroborating the view that the free cysteines might also contribute to sulfide binding in B1 hemoglobin chain of deep-sea tubeworms (Li et al., 2019).

**Figure 5.**
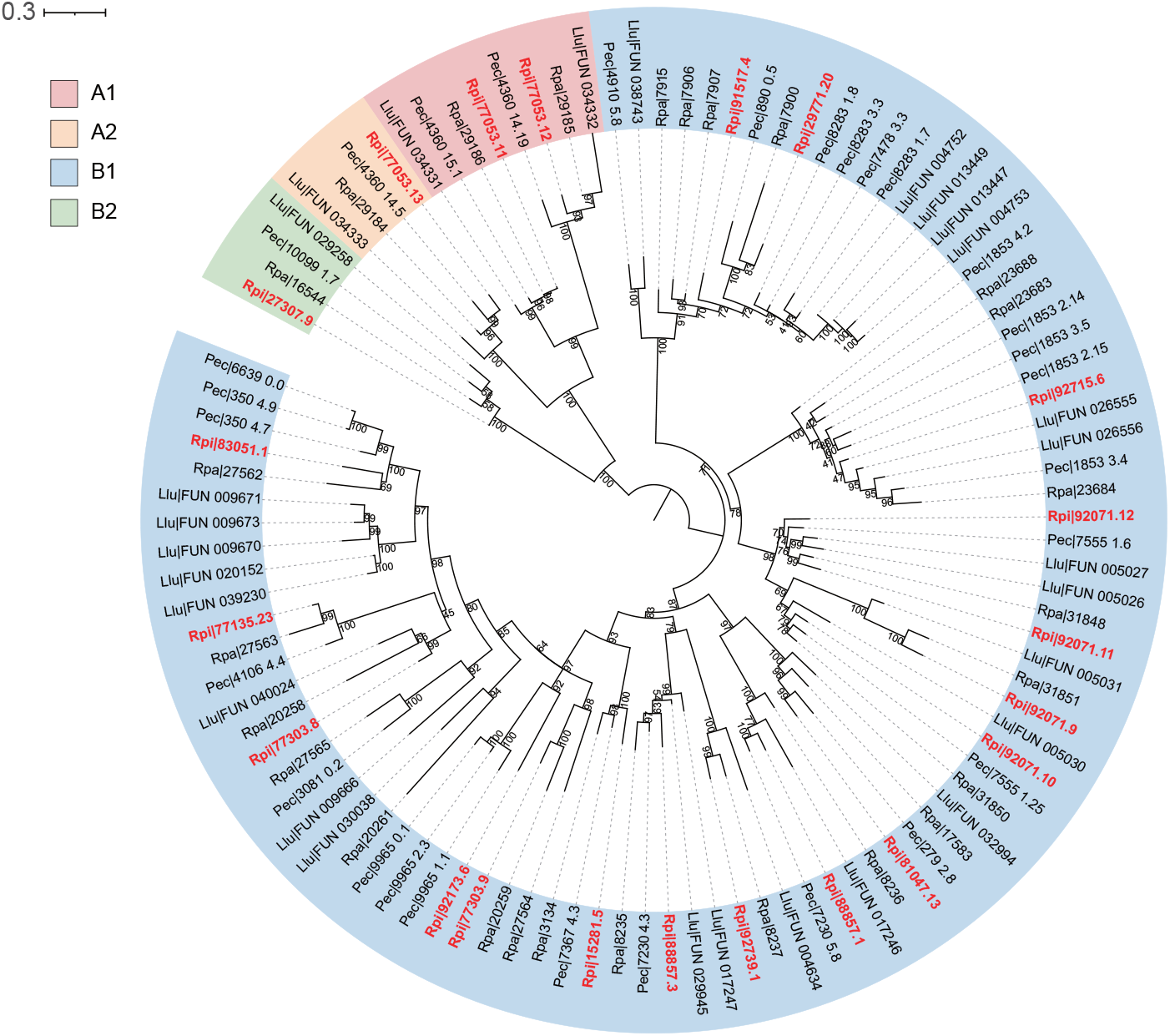
Gene tree of Hb subunits A1, A2, B1, and B2 from four vestimentiferan tubeworms. The values near the nodes are ultrafast bootstrap (UFBoot) values. *R. piscesae* sequences are labeled red.

Recent reports revealed that most enzymes related to amino acid biosynthesis were lost in *L. luymesi* and *R. pachyptila* (Li et al., 2019; de Oliveira et al., 2022). To gain better insight into nutrient dependence of endosymbionts in vestimentiferans, we identified key enzymes involved in amino acid biosynthesis in the genomes of tubeworms (**Fig. 6**). All four tubeworms (*L. luymesi* and *R. pachyptila*, *R. piscesae* and *P. echinospica*) lack most key enzymes related to amino acid biosynthesis, corroborating the view that vestimentiferan tubeworms mainly relied on endosymbionts for synthesizing amino acids (Li et al., 2019).

**Figure 6.**
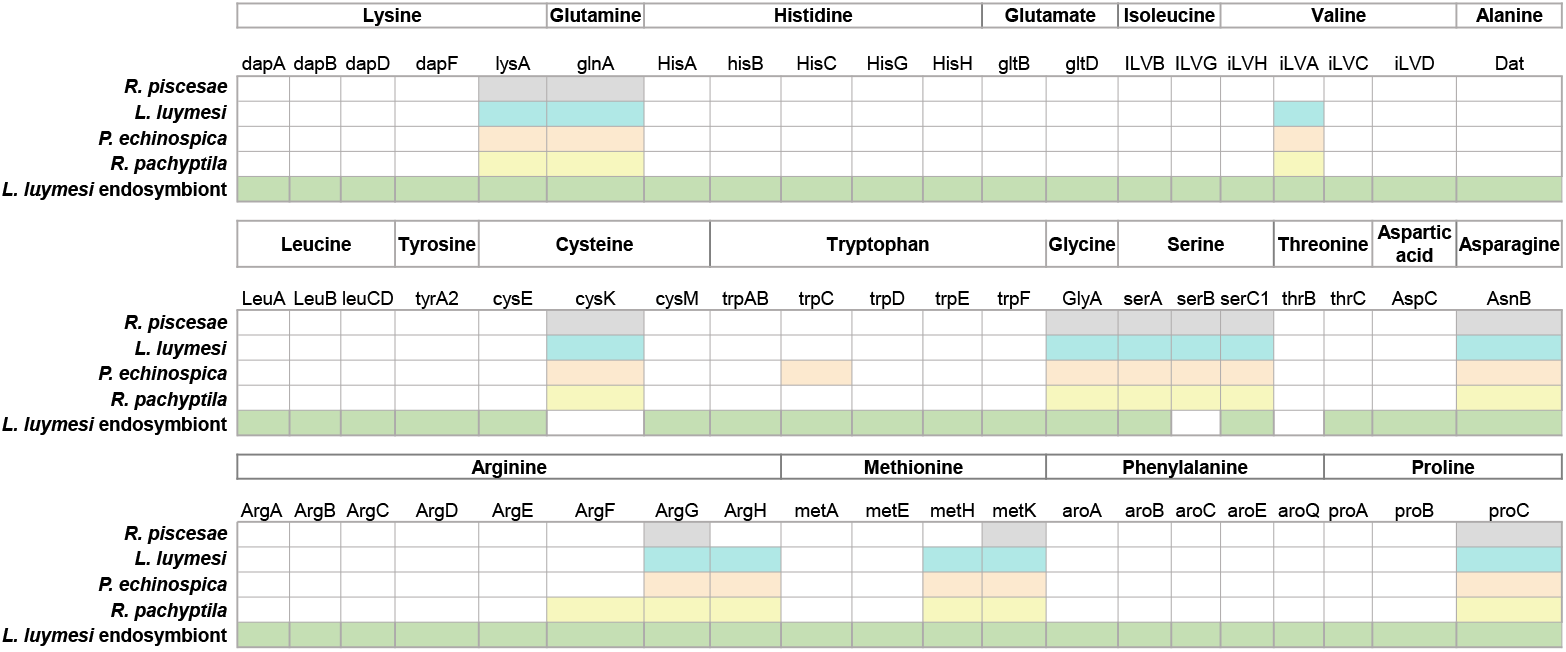
Amino acid biosynthesis genes. Most key genes associated with amino acid biosynthesis are missing in the genomes of *L. luymesi, P. echinospica, R. pachyptila,* and *R. piscesae.* These genes are presented in the genome of *L. luymesi* symbionts.

The expansion of gene families is considered as a major driver of adaptation and speciation (Sharpton et al., 2009). Thus, we performed gene family expansion and contraction analysis with 4 vestimentiferan tubeworms and 8 other lophotrochozoans (**Fig. 4B**). In total, 10 gene families were significantly expanded in the genomes of all four tubeworms compared to other 8 lophotrochozoans (*P* < 0.05) (**Supplementary Table S10**). Gene ontology analysis revealed the expanded gene families were involved in the process of chitin binding and innate immunity. Furthermore, 18 gene families were significantly expanded in the genomes of two cold-seep tubeworms. The expanded gene families were involved in the process of DNA repair, innate immunity, and protein stability (**Supplementary Table S11**).

In addition to expanded and contracted gene families, we identified positively selected genes (PSGs) in the genome of *R. piscesae.* Compared with other 11 lophotrochozoan species, 9 PSGs were identified in *R. piscesae* (**Supplementary Table S12**). Interestingly, four genes (alkB homolog 2, alpha-ketoglutarate-dependent dioxygenase, *ALKBH2;* Derlin-1, *DERL1;* Ras-related and estrogen-regulated growth inhibitor, *RERG;* AN1-type zinc finger protein 2B, *ZFAND2B*) involved in cell proliferation were subjected to positive selection in *R. piscesae*. Two of these genes (*ALKBH2* and *DERL1*) promote cell proliferation. ALKBH2 is responsible for protecting the genome from 1-meA damage by repairing the damage in double-stranded DNA(Aas et al., 2003). *ALKBH2* promotes cell proliferation and is overexpressed in several types of tumor cells (Wilson et al., 2018). DERL1 participates in endoplasmic reticulum (ER)- associated degradation response and unfolded protein response (UPR) (Eshraghi et al., 2014). *DERL1* is responsible for cell proliferation and promotes the progression of several types of cancers (Dong et al., 2013). Interestingly, two other genes *(RERG* and *ZFAND2B*) inhibit cell proliferation. Overexpression of *RERG*, a member of the RAS superfamily of GTPase, inhibits cell proliferation and tumor formation (Finlin et al., 2001; Ho et al., 2017). ZFAND2B reduces the abundance of IGF1R, a kinase that activates cell proliferation, in a proteasome-dependent manner (Osorio et al., 2016). These results suggest that, besides apoptosis, the regulation of cell proliferation also contributes to the variation of growth rates in *R. piscesae*.

## Discussion

In many hydrothermal vent and cold seep ecosystems, vestimentiferan tubeworms are among the dominant megafauna at habitats where hydrogen sulfide is present (Cavanaugh et al., 1981; Tunnicliffe, 1992). These deep-sea tubeworms occupy a broad environmental gradient from areas where sulfide-rich fluids emanate as relatively vigorous and continuous diffuse vent flow to areas where vent fluids seep slowly from the seafloor (Urcuyo et al., 2007). Seep-dwelling tubeworms grow slowly as they live in environments where exposure to seep fluid is very low (Julian et al., 1999; Fisher et al., 2008). The lifespan of most vent-dwelling tubeworms is relatively short due to the habitats are short-lived. They thrive in strong and continuous diffuse hydrothermal flow and dies when vent flow subsides or sulfide levels decrease due to biotic or abiotic factors (Fisher et al., 1988; Shank et al., 1998). Great variations in lifespan and morphology were found among different morphotypes of *R. piscesae*, which is endemic to the highly heterogeneous chemical environments of hydrothermal vents on the Juan de Fuca Ridge. The “Short-fat” morphotype of *R. piscesae* that live around vigorous vents with high concentrations of sulfide grow rapidly, while the “long-skinny” morphotype inhabit around diffuse flow with low concentrations of sulfide have much slower growth rates (Southward et al., 1995; Carney et al., 2007). Unlike the “short-fat” morphotype, the tube diameter of “long-skinny” morphotype reduces from the anterior to posterior. As the concentration of sulfide is low in the diffuse vent flow, thin and transparent end facilitates the “long-skinny” morphotype to acquire sulfide by the posterior tube (Urcuyo et al., 2003). Thus, *R. piscesae* exhibit a greater tolerance to varying physicochemical conditions than any other known vestimentiferans (Tivey et al., 1999; Bright and Lallier, 2010). However, the evolutionary history and underlying genetic mechanisms of adaptation in *R. piscesae* are remained to be investigated.

Here we assembled and annotated a high-quality reference sequence of *R. piscesae*. Phylogenomic analysis was performed among 4 vestimentiferan tubeworms and 11 metazoans using single-copy orthologs (**Fig. 1, Supplementary Fig. S2**). *Ridgeia pachyptila* and *R. piscesae*, the two tubeworm species that are endemic to hydrothermal vents of the eastern Pacific, formed a clade in the phylogenetic tree. Additionally, *Lamellibrachia luymesi* from cold seep appeared sister to three other deep-sea tubeworms, corroborating the view that this species derived early from other tubeworm (Black et al., 1997). *Ridgeia piscesae* and *R. pachyptila* are endemic to the GFE ridge systems and the EPR, respectively. The divergence time between these two vent-dwelling species was estimated to be approximately 33.7 Ma. This result suggests that *R. piscesae* and *R. pachyptila* started to diverge after the formation of GFE and EPR between 28.5-35 Ma.

It was proposed that natural selection and adaptive process shaped genome size evolution (Gregory, 2001). Previous analyses revealed that genome sizes are correlated with several phenotypic traits, including cell size, rates of metabolism and growth (Cavalier-Smith, 1982; Vinogradov, 1995; Wyngaard et al., 2005; Wright et al., 2014). A negative correlation between genome size and growth rate was identified in several species, as organisms with smaller genomes might undergo more rapid replication time of their genome (Wyngaard et al., 2005; Tenaillon et al., 2016). To investigate the genomic basis of their high growth rates, we studied the factors which might contribute to the evolution of genome sizes in four vestimentiferan tubeworms. Firstly, we studied the relationship between repeat content and genome size. A strong positive correlation (R^2^ = 0.98, *P* = 0.0052) between repeat content and genome size was identified in these tubeworm species (**Fig. 2B**), corroborating the view that repeat content contributes to the variation of genome sizes in tubeworm species (Sun et al., 2021). Secondly, we evaluated the number and size of genes in tubeworm genomes. *Lamellibrachia luymesi* genome has the most annotated gene models (38,998), following by *R. pachyptila* (25,984), *R. piscesae* (24,096), and *P. echinospica* (22,642). Average length of introns in the genome of *R. piscesae* is longer than introns of other three species. In addition, two vent-dwelling tubeworms with smaller genome sizes have higher ratios of intron / exon length than the seep-dwelling tubeworms. These results suggest that the variation of genome sizes in tubeworms is not attributed to gene number and intron length. Lastly, we performed gene family expansion and contraction analysis. Lineage-specific gene families of *R. piscesae* (601) and *R. pachyptila* (279) are much less than the ones of *L. luymesi* (1181) and *P. echinospica* (1045) (**Fig. 4A**). In addition, gene-family expansion and contraction analysis revealed that the numbers of expanded gene families were substantially less than contracted gene families in the two vent-dwelling tubeworms (**Fig. 4B**). These results indicate that, besides *R. pachyptila*, *R. piscesae* experienced reductive evolution. Taken together, these results indicate that the high growth rates of vent-dwelling tubeworms might derive from small genome size and protein family. The small genome sizes of these tubeworms are attributed to the repeat content but not the number of genes and intron sizes.

Previous immunohistochemical and ultrastructural cell cycle analyses revealed that cell proliferation activities of *L. luymesi* and *R. pachyptila* are as high as in tumor. The balanced activities of proliferation and apoptosis in the epidermis lead to the slow growth in *L. luymesi* from cold seeps, while apoptosis is substantially downregulated in this tissue of *R. pachyptila* that maintains a high growth rate (Pflugfelder et al., 2009). Unlike other vestimentiferan tubeworms, growth rates vary greatly among different individuals of *R. piscesae* (Tunnicliffe et al., 1997; Sarrazin et al., 1998). Four genes involved in the regulation of cell proliferation were identified to be positively selected in *R. piscesae*. Interestingly, two of these genes promote cell proliferation, whereas two other genes inhibit cell proliferation. This result indicates that both cell proliferation and apoptosis involve in the regulation of growth in *R. piscesae.* Furthermore, up-regulation and downregulation of cell proliferation might play important role in the variation of growth rate in this species.

## Materials and Methods

### Sampling and sequencing

The samples of *R. piscesae* were obtained during *Alvin* dive 4243 from the deep-sea hydrothermal vent at Cathedral vent, Main Endeavor Field of the Juan de Fuca Ridge (47° 56’ N, 129° 05’ W, 2,181 m depth) on August 9, 2006. Genomic DNA (gDNA) was extracted from muscle of the specimen using a standard phenol/chloroform extraction protocol and broken into random fragments for whole-genome shotgun (WGS) sequencing. Agarose gel electrophoresis was used to check the quality of the gDNA, and Qubit system was used to quantify the gDNA. Short-insert paired-end libraries (180 bp, 300 bp and 500 bp) were prepared using the NEBNext Ultra DNA Library Prep Kit for Illumina (NEB) according to the standard protocol, respectively. Large-insert mate-pair libraries (2 kb, 5 kb, 10 kb and 15 kb) were prepared following the Cre-lox recombination-based protocol (Van Nieuwerburgh et al., 2012). All DNA libraries were subjected to paired-end sequencing on the Illumina Hiseq 2000 platform (Illumina). Muscle samples were also collected for constructing RNA sequencing (RNA-seq) library. Total RNA was extracted with TRIzol reagent (Molecular Research Center). Paired-end library for RNA-seq was constructed using the Paired-End Sample Preparation Kit (Illumina) and sequenced on the Illumina Hiseq 2000 platform (Illumina).

### Genome assembly

NGS QC toolkit (v2.1) (Patel and Jain, 2012) was used to evaluate the quality of raw sequencing reads and filter high-quality reads. High-quality reads were obtained by filtering out the following types of reads: (1) reads with >= 10% unidentified nucleotides (N); (2) reads with adaptor contamination; (3) reads with >= 20% bases having Phred quality score <= 5; (4) duplicated reads generated by PCR amplification during the library construction process.

The size and heterozygosity of *R. piscesae* genome were estimated using the high-quality short-insert paired-end reads (180 bp) by *k*-mer frequency-distribution method. The number of *k*-mers and the peak depth of *k*-mer sizes at 19 was obtained using GenomeScope2 (v1.0.0) (Ranallo-Benavidez et al., 2020). Due to the high heterozygosity of *R. piscesae* genome, a modified version of SOAPdenovo (Wang et al., 2017) was implemented for genome assembly. In brief, all short-insert paired-end reads were applied for contig assembly. Depth of coverage was obtained for each contig using SOAPdenovo with the parameters ‘-e 1 -M 0 -R’, and the contigs with depth less than 60 were identified as heterozygous contigs. All WGS reads were aligned to the heterozygous contigs using SOAPdenovo. And links were generated between heterozygous contigs when supported by a minimum of three read pairs. Heterozygous contigs were clustered into bubble clusters based on the orientation and distance between heterozygous contigs. If two contigs represented two potential haplotypes in a bubble structure, the longer one was retained to ensure the integrity of contig assembly.

To scaffolding the contigs, all short-insert paired-end and long-insert mate-pair reads were realigned onto the contig sequences using SOAPdenovo. Duplicated contigs that had high depth of coverage and conflicting connections to the unique contigs were masked during scaffolding. A hierarchical assembly strategy was used to construct contigs into primary scaffolds by adding the ascending insert size reads gradually. Finally, all short-insert reads were realigned onto the scaffold sequences to fill the gaps with the GapCloser program implemented in SOAPdenovo (Luo et al., 2012).

### Genome quality assessment

To assess the completeness of the *R. piscesae* genome, high-quality short-insert paired-end reads were mapped to the genome assembly using Burrows-Wheeler Aligner (BWA) (v0.7.17) (Li and Durbin, 2009) with parameters of ‘-o 1 -e 5 -t 8 -n 15’. In addition, all RNA-seq reads were *de novo* assembled using Trinity (v2.9) (Grabherr et al., 2011). The Trinity assembled sequences (Unigenes) with length >= 500 bp were mapped to the *R. piscesae* genome using BLAT (v35.1) (Kent, 2002) with default parameters and an identify cutoff of 90%. The completeness of the assembly was also evaluated using benchmarking universal single-copy orthologs (BUSCO) (v3.1.0) (Simao et al., 2015) with 978 metazoa single-copy orthologous genes (obd10).

### Genome annotation

Tandem repeats in the genome were predicted using the program Tandem Repeats Finder (TRF) (v4.09) (Benson, 1999) with default parameters. Transposable elements (TEs) were identified using the homology-based and *de novo* prediction approaches. For homology-based prediction, RepeatMasker (v4.1.0) (http://www.repeatmasker.org/) were conducted to identify repeat sequences against the Repbase library. For *de novo* prediction, RepeatModeler (v2.0.1) (http://repeatmasker.org/RepeatModeler.html), LTR-Finder (v1.0.7) (Xu and Wang, 2007), RepeatScout (v1.0.5) (Price et al., 2005) and Piler (v1.0) (Edgar and Myers, 2005) were used to construct *de novo* repeat libraries. RepeatMasker (v4.1.0) was run against these libraries to search repeat elements.

Protein-coding genes in *R. piscesae* genome were predicted with three approaches: homology-based prediction, *ab initio* prediction and RNA-seq-based prediction. Protein-coding sequences of *Lottia gigantea, Helobdella robusta, Capitella teleta, Schistosoma mansoni, Caenorhabditis elegans, Anopheles gambiae, Drosophila melanogaster* and *Homo sapiens* were aligned to the *R. piscesae* genome using tblastn with a cut off E-value of 1e-5. GeneWise (v2.4) (Birney et al., 2004) was employed to predict gene models. For *ab initio* prediction, Augustus (v3.3.2) (Stanke and Morgenstern, 2005), Genscan (Aggarwal and Ramaswamy, 2002), Geneid (v1.3) (Parra et al., 2000), GlimmerHMM (v3.0.4) (Majoros et al., 2004) and SNAP (Korf, 2004) were used to predict genes on the repeat-masked genome. For RNA-seq-based prediction, Trinity (v2.9) generated sequences (Unigenes) were aligned against the genome assembly with BLAT (v35.1) (identify >= 0.95 and align rate >= 0.95) (Kent, 2002). In addition, the RNA-seq reads from were aligned to the *R. piscesae* genome using Tophat (v2.1.1) (Trapnell et al., 2009). And gene structures were predicted using Cufflinks (v2.2.1) (Trapnell et al., 2010). EvidenceModeler (EVM) (v1.1.1) (Haas et al., 2008) was used to integrate all gene models derived from these three approaches into a non-redundant gene set.

Functional annotation was performed using BLASTP searches against SwissProt and TrEMBL databases (Bairoch and Apweiler, 2000) with a E-value cut-off of 1e-5. In addition, InterProScan (v5.4.0) (Mulder and Apweiler, 2007) was used to screen proteins against five databases (Pfam, PRINTS, PROSITE, ProDom and SMART) to determine protein domains and motifs. Gene Ontology (GO) annotation of each gene was retrieved from the corresponding InterPro entry. In addition, KEGG annotation was performed using GhostKOALA (Kanehisa et al., 2016).

### Phylogenomic analysis

Protein sequences of 14 metazoan species *(Adineta vaga, Echinococcus multilocularis, Aplysia californica, Lottia gigantea, Octopus bimaculoide, Phoronis australis, Lingula anatina, Notospermus geniculatus, Capitella teleta, Helobdella robusta, Eisenia Andrei, Riftia pachyptila*, *Paraescarpia echinospica*, *Lamellibrachia lumysi*) were downloaded for gene family cluster analysis (**Supplementary Table S8**). The longest transcripts of each gene (more than 30 amino acids) were retained. All-to-all BLASTP was used to identify the similarities between retained protein sequences of these 14 metazoan species and *R. piscesae* (E-value threshold: 1e-7). OrthoFinder (v2.2.7) (Emms and Kelly, 2019) was used to identify and cluster gene families among 15 species with default parameters. Gene clusters with >100 gene copies in one or more species were removed. Protein sequences of all single-copy gene families were retrieved and aligned using MAFFT (v7.271) (Katoh et al., 2002). The alignments were trimmed using TrimAI (v1.2) (Capella-Gutierrez et al., 2009). The phylogenetic tree was reconstructed with the trimmed alignments using FastTree2 (v2.1.11) (Price et al., 2010) with *Adineta vaga* as outgroup.

To estimate the divergent time, the trimmed alignments of single-copy orthologs among the 15 metazoan species were concatenated using PhyloSuite (v1.2.2) (Zhang et al., 2020). MCMCtree module of the PAML package (v4.9) (Yang, 2007) was used to estimate the divergent time with the concatenated alignment. The species tree of the 15 metazoan species was used as a guide tree, and the analysis was calibrated with the divergent time obtained from TimeTree database (minimum = 470.2 million years and soft maximum = 531.5 million years between *P. australis* and *L. anatina)* (Kumar et al., 2017) and previous analyses (minimum = 470.2 million years and soft maximum = 531.5 million years between *A. californica* and *L. gigantea;* minimum = 532 million years and soft maximum = 549 million years for the first appearance of Mollusca; minimum = 476.3 million years and soft maximum = 550.9 million years for the appearance of capitellid-leech clade; minimum = 550.25 million years and soft maximum = 636.1 million years for the first appearance of Lophotrochozoa and Ecdysoa) (Donoghue et al., 2009; Benton et al., 2015; dos Reis et al., 2015).

### Gene family expansion and contraction analysis

r8s (v1.7) was applied to obtain the ultrametric tree of 12 lophotrochozoan species *(C. teleta, H. robusta, E. andrei, L. gigantea, A. californica, N. geniculatus, A. californica, P. australis*, *R. pachyptila*, *P. echinospica*, *L. lumysi*, *R. piscesae*), which is calibrated with the divergent time between *C. teleta* and *L. gigantea* (688 mya) obtained from TimeTree database. CAFÉ (v5) (De Bie et al., 2006) was applied to determine the significance of gene-family expansion and contraction among 12 lophotrochozoan species based on the ultrametric tree and the gene clusters determined by OrthoFinder (v2.2.7). Gene families that were significantly expanded in each of four tubeworm species *(R. pachyptila, P. echinospica, L. lumysi, R. piscesae*) (*P* < 0.05) were annotated using PANTHER (v16.0) with the PANTHER HMM scoring tool (pantherScore2.pl) (Mi et al., 2017).

### Homeobox gene analysis

Homeodomain sequences, which were retrieved from HomeoDB database(Zhong and Holland, 2011), were aligned to *R. piscesae* genome assembly using tbalstn. Sequences of the candidate homeobox genes were extracted based on the alignment results. The extracted sequences were aligned against NCBI NR and HomeoDB database to classify the homeobox genes.

### Hemoglobin gene family analysis

Protein sequences of hemoglobin A1, A2, B1, B2 chains of four tubeworm species were obtained with reference references using DIAMOND BLASTP (Buchfink et al., 2021) with a E-value cut-off of 1e-5. The sequences were annotated in NCBI NR database using BLASTP. And protein domains in these sequences were annotated by Pfamscan against Pfam-A.hmm database (Finn et al., 2014). Sequences that have almost full length protein domains were aligned using MAFFT (v7.271) (Katoh and Standley, 2013). The alignments were trimmed using TrimAI (v1.2) (Capella-Gutierrez et al., 2009). The phylogenetic tree was reconstructed with the trimmed alignments using a maximum-likelihood method implemented in IQ-TREE2 (v2.1.2) (Minh et al., 2020). The best-fit substitution model was selected by using ModelFinder algorithm (Kalyaanamoorthy et al., 2017). Branch supports were assessed using the ultrafast bootstrap (UFBoot) approach with 1,000 replicates (Hoang et al., 2018).

### Identification of positively selected genes (PSGs)

We identified PSGs in the *R. piscesae* genome within single-copy orthologs among 12 lophotrochozoan species that were identified in gene-family expansion and contraction analysis. Protein sequences of all single-copy gene families were retrieved and aligned using MAFFT (v7.271) (Katoh et al., 2002). Phylogenetic tree of each family was reconstructed using IQ-TREE2 (v2.1.2) (Minh et al., 2020). PSGs were identified based on the phylogenetic trees using HyPhy (v2.5.30) with the adaptive Branch-Site Random Effects Likelihood (aBSREL) model (Pond et al., 2020).

## Data Availability

Raw reads and genome assembly are accessible in NCBI under BioProject number PRJNA826206. Assembled genome sequences are accessible under Whole Genome Shotgun project number JALOCR000000000. Raw reads and genome assembly are also available at the CNGB Sequence Archive (CNSA) of China National GeneBank DataBase (CNGBdb) with accession number CNP0002911.

## Acknowledgements

This work was supported by National Key R&D Program of China (2018YFC0310702), the Projects under Major State Basic Research Development Program of China (973 Program) (2015CB755906), National Natural Science Foundation of China (No. 31900309), GuangDong Basic and Applied Basic Research Foundation (No. 2019A1515011644), Innovation Group Project of Southern Marine Science and Engineering Guangdong Laboratory (Zhuhai) (No. 311021006), and Project 2018N2001 from Department of Fujian Science and Technology and Program for Innovative Research Team in Science and Technology in Fujian Province University. The funders had no role in study design, data collection and analysis, decision to publish, or preparation of the paper. We gratefully acknowledge the National Supercomputing Center in Guangzhou for provision of computational resources.

## Author contributions

J.M.C, C.J.G and J.G.H. conceived of the project and designed research; L.R., M.L., Z.L, Y.W. assembled and annotated the genome; M.W, J.H., L.Z., H.S., M.C., Y.J., F.Y., and R.Z. performed the evolutionary analyses; M.W., J.M.C., C.J.G, and J.G.H. wrote the paper with contribution from all authors.

## Ethics declarations

### Animal and human rights statement

The tubeworm used in our study is an invertebrate, so the approval according to the regulations on the use of tubeworm is unnecessary.

