## Supplementary Materials for "The genome of a vestimentiferan tubeworm (*Ridgeia piscesae*) provides insights into its adaptation to a deep-sea environment"

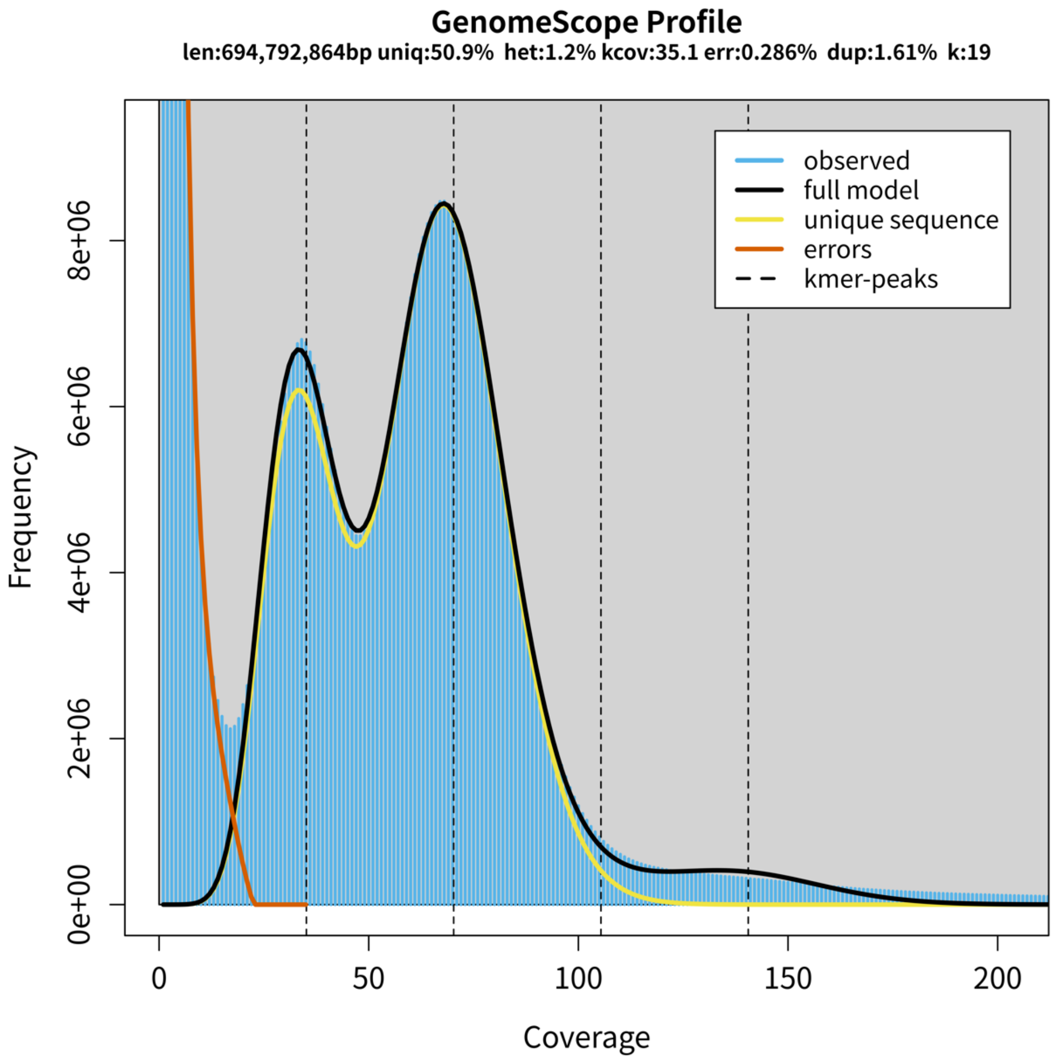


**Supplementary Figure S1. Distribution of 19-mer frequency in *Ridgeia piscesae* genome.** The short-insert paired-end reads (180 bp) were used to generate the 19-mer frequency curve. The heterozygous rate and the genome size were determined based on the *k*-mer distribution.

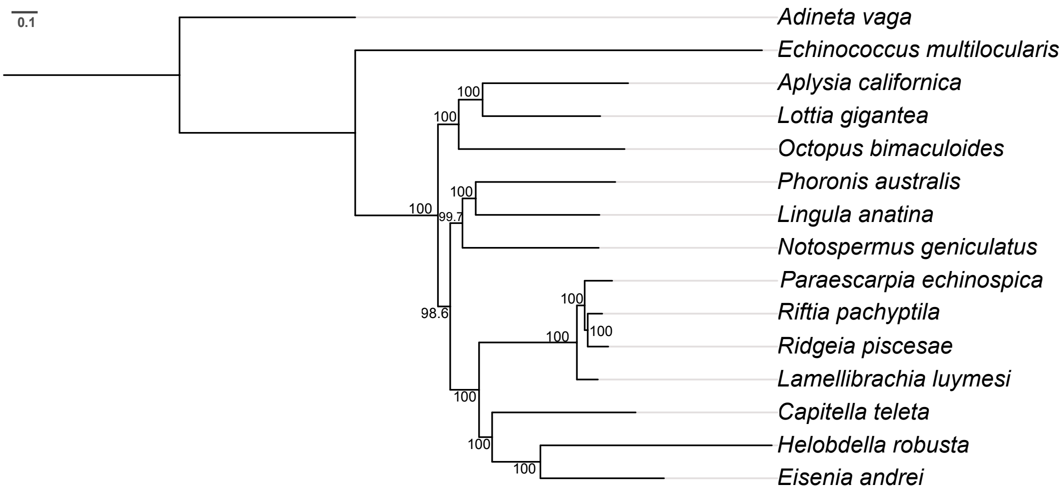


**Supplementary Figure S2. The phylogenetic tree of *R. piscesae* and 14 other lophotrochozoans.** The tree was reconstructed with single-copy orthologs using a maximum likelihood approach. The ultrafast bootstrap (UFBoots) value is listed above each of the nodes.

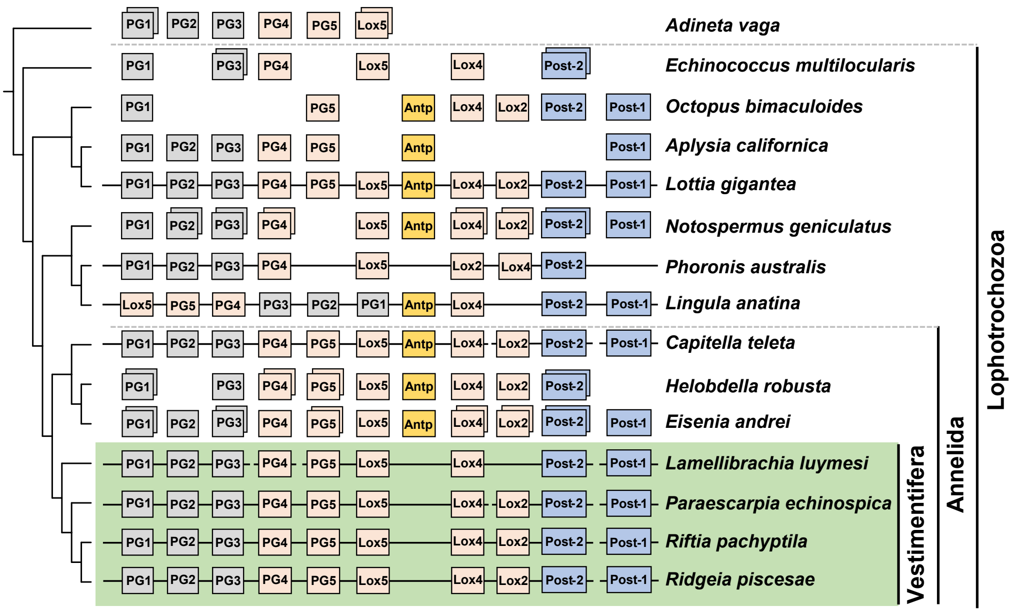


**Supplementary Figure S3. Genomic organization of *Hox* gene clusters in 4 vestimentiferan tubeworms and 11 other metazoans.** *Hox* genes are indicated as rectangles. The orientations of genes are indicated by arrows below the genes. The gene composition and orientation of *Hox* clusters are consistent between two vent-dwelling tubeworms (*R. pachyptila* and *R. piscesae*), but slightly different between vent- and seep-dwelling tubeworms.

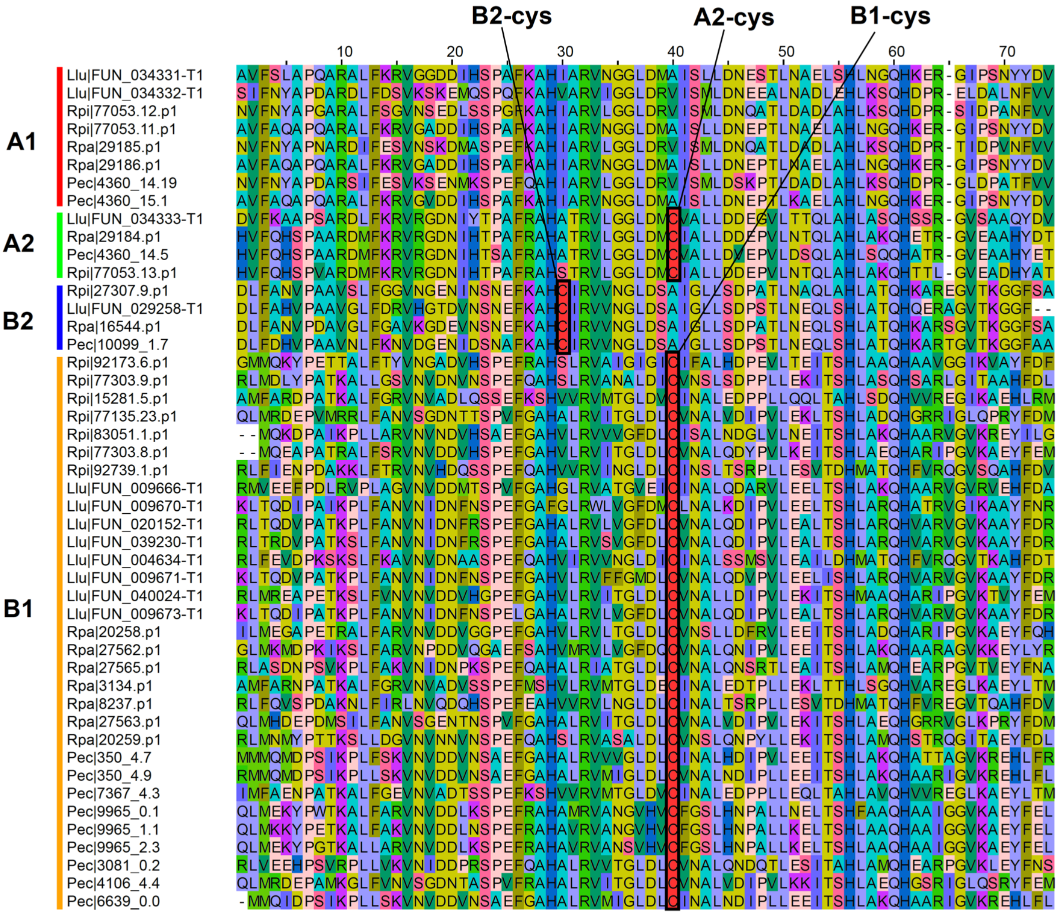


**Supplementary Figure S4. Alignment of hemoglobins in four tubeworms.** Each tubeworm has two copies of A1 chain, one copy of A2 chain, and one copy of B2 chain in hemoglobins of tubeworms. A group of B1 chain in hemoglobin were found in each of four species. Free cysteine was found in A2, B2, and B1 chains in hemoglobin.

**Supplementary Table S1.** Statistics of the genome sequencing data of *Ridgeia piscesae*

| Pair-end libraries | Insert size | Raw data (Gb) | Read length (bp) | Sequence coverage (X) † |
|---|---|---|---|---|
| Illumina data | 180 bp | 87.13 | 100 | 125.40 |
| | 300 bp | 40.62 | | 58.46 |
| | 500 bp | 38.74 | | 55.76 |
| | 2 kb | 18.73 | | 26.96 |
| | 5 kb | 20.76 | | 29.88 |
| | 10 kb | 31.85 | | 45.84 |
| | 15 kb | 9.91 | | 14.26 |
| Total | - | 247.74 | - | 356.56 |

†Sequencing coverage was calculated with the clean data and the estimated genome size of 694.79 Mb by *k*-mer analysis

**Supplementary Table S2.** Statistic of the *R. piscesae* genome assembly

| Sample ID | Contig | | Scaffold † | |
|---|---|---|---|---|
| | **number** | **Length** | **number** | **Scaffold(bp)** |
| Total | 113,966 | 529,732,904 | 29,336 | 574,959,875 |
| Max | - | 174,253 | - | 2,042,129 |
| Number>=100 | 113,736 | - | 29,336 | - |
| Number>=2000 | 57,711 | - | 7,801 | - |
| N50 | 12,867 | 10,417 | 750 | 230,234 |
| N60 | 18,770 | 7,765 | 1,035 | 178,368 |
| N70 | 26,816 | 5,575 | 1,412 | 128,609 |
| N80 | 38,421 | 3,718 | 1,967 | 81,082 |
| N90 | 57,494 | 2,014 | 3,079 | 30,247 |

†Only scaffolds with length >= 200 bp were counted.

**Supplementary Table S3.** Assessment of genome coverage rate based on short-insert paired-end reads remapping analysis

| | | **Percentage (%)** |
|---|---|---|
| Reads | Mapping rate (%) | 87.4 |
| Genome | Average sequencing depth | 71.98 |
| | Coverage (%) | 99.74 |
| | Coverage at least 4X (%) | 98.61 |
| | Coverage at least 10X (%) | 96.58 |
| | Coverage at least 20X (%) | 92.92 |

**Supplementary Table S4.** Assessment of gene coverage rate using Trinity assembled sequences (Unigenes)

| Dataset | Number | Total length (bp) | Sequences covered by assembly (%) | With > 90% sequence in one scaffold | | With > 50% sequence in one scaffold | |
|---|---|---|---|---|---|---|---|
| | | | | Number | Percent | Number | Percent |
| > 0bp | 18,529 | 21,132,160 | 99.63 | 16,543 | 89.28 | 18,207 | 98.26 |
| > 200bp | 18,529 | 21,132,160 | 99.63 | 16,543 | 89.28 | 18,207 | 98.26 |
| > 500bp | 18,529 | 21,132,160 | 99.63 | 16,543 | 89.28 | 18,207 | 98.26 |
| > 1000bp | 7,325 | 13,371,531 | 99.89 | 6,415 | 87.58 | 7,178 | 97.99 |
| > 2000bp | 2,118 | 6,230,088 | 100.00 | 1,903 | 89.85 | 2,069 | 97.69 |
| > 5000bp | 87 | 554,752 | 100.00 | 74 | 85.06 | 83 | 95.40 |

**Supplementary Table S5** BUSCO evaluation of *R. piscesae* genome assembly

| | *R. piscesae* |
|---|---|
| Complete BUSCOs | 884 |
| Complete and single-copy BUSCOs | 878 |
| Complete and duplicated BUSCOs | 6 |
| Fragmented BUSCOs | 27 |
| Missing BUSCOs | 43 |
| Total BUSCO groups searched | 954 |

**Supplementary Table S6** Summary of annotated repeats in *R. piscesae* genome

| | Number | Length（bp） | Percentage（%） |
|---|---|---|---|
| **Retroelements** | **203,600** | **56,094,618** | **9.76** |
| SINEs: | 26,182 | 5,142,860 | 0.89 |
| Penelope | 5,103 | 888,385 | 0.15 |
| LINEs: | 164,545 | 46,431,336 | 8.08 |
| CRE/SLACS | 313 | 32329 | 0.01 |
| L2/CR1/Rex | 106,412 | 29,637,674 | 5.15 |
| R1/LOA/Jockey | 61 | 10,440 | 0.00 |
| R2/R4/NeSL | 394 | 118,283 | 0.02 |
| RTE/Bov-B | 39,673 | 12,731,947 | 2.21 |
| L1/CIN4 | 3,031 | 249,962 | 0.04 |
| LTR: | 12,873 | 4,520,422 | 0.79 |
| BEL/Pao | 33 | 6,420 | 0.00 |
| Ty1/Copia | 8 | 1,185 | 0.00 |
| Gypsy/DIRS1 | 8,533 | 3,448,083 | 0.60 |
| Retroviral | 456 | 24,641 | 0.00 |
| **DNA transposons：** | **35,533** | **10,993,251** | **1.91** |
| hobo-Activator | 14,208 | 3,067,637 | 0.53 |
| Tc1-IS630-Pogo | 2976 | 1,100,786 | 0.19 |
| PiggyBac | 2 | 147 | 0.00 |
| Tourist/Harbinger | 1,511 | 760,090 | 0.13 |
| Other (Mirage,P-element,Transib) | 905 | 214,308 | 0.04 |
| **Rolling circles** | **1,933** | **211,701** | **0.04** |
| **Unclassified:** | **546,050** | **106,343,896** | **18.50** |
| **Small RNA:** | **1,290** | **152,996** | **0.03** |
| **Satellites:** | **547** | **175,801** | **0.03** |
| **Simple repeats:** | **282,603** | **24,492,855** | **4.26** |
| **Low complexity** | **9,714** | **828,375** | **0.14** |

**Supplementary Table S7.** Statistics of functional annotated gene models in the genome of *R. piscesae*

| | **Number** | **Percentage (%)** |
|---|---|---|
| InterPro | 16,309 | 67.68 |
| GO | 16,741 | 69.48 |
| Pfam | 16,068 | 66.68 |
| Swissprot | 17,849 | 74.07 |
| TrEMBL | 21,721 | 90.14 |
| KEGG | 17,022 | 70.64 |
| **Annotated** | 23,021 | 95.54 |
| **Unannotated** | 1,075 | 4.46 |
| Total | 24,096 | - |

**Supplementary Table S8** Information of genomes used to perform phylogenomic analysis

| Species name | NCBI ID | Reference |
|---|---|---|
| *Lamellibrachia lumyesi* | GCA_009193005.1 | (Li et al., 2019) |
| *Paraescarpia echinospica* | GCA_020002185.1 | (Sun et al., 2021) |
| *Ridgeia piscesae* | - | This study |
| *Riftia pachyptila* | - | (de Oliveira et al., 2022) |
| *Capitella teleta* | GCA_000328365 | (Simakov et al., 2013) |
| *Helobdella robusta* | GCA_000326865.1 | (Simakov et al., 2013) |
| *Eisenia andrei* | - | (Shao et al., 2020) |
| *Lingula anatina* | GCA_001039355.2 | (Luo et al., 2015) |
| *Aplysia californica* | GCA_000002075.2 | - |
| *Octopus bimaculoides* | GCA_001194135 | (Albertin et al., 2015) |
| *Notospermus geniculatus* | GCA_002633025.1 | (Luo et al., 2018) |
| *Lottia gigantea* | GCA_000327385 | (Simakov et al., 2013) |
| *Phoronis australis* | GCA_002633005.1 | (Luo et al., 2018) |
| *Adineta vaga* | GCA_021613535.1 | (Flot et al., 2013) |
| *Echinococcus multilocularis* | GCA_000469725.3 | (Tsai et al., 2013) |

**Supplementary Table S9** Exon and intron lengths of genes in four Vestimentiferan tubeworms

| Species | Genome size (Mb) | Total Exon Length (bp) | Total Intron Length (bp) | Mean Exon Length (bp) | Mean Intron Length (bp) | Intron / Exon length |
|---|---|---|---|---|---|---|
| ***Lamellibrachia luymesi*** | 687.7 | 42,059,392 | 220,083,629 | 204.18 | 1332.65 | 6.53 |
| ***Paraescarpia echinospica*** | 1090.9 | 57,384,225 | 204,708,487 | 375.12 | 1570.63 | 4.19 |
| ***Riftia pachtypila*** | 560.7 | 46,707,225 | 264,839,561 | 229.08 | 1531.29 | 6.68 |
| ***Ridgeia piscesae*** | 574.9 | 36,771,027 | 234,465,512 | 223.80 | 1672.26 | 7.47 |

**Supplementary Table S10** Gene families were significantly expanded in the genomes of all four tubeworms

| | | **P. echinospica** | **L. luymesi** | **R. piscesae** | **R. pachyptila** | **C. teleta** | **H. robusta** | **E. andrei** | **L. gigantea** | **A. californica** | **O. bimaculoides** | **P. australis** | **N. geniculatus** |
|---|---|---|---|---|---|---|---|---|---|---|---|---|---|
| **OG0000338** | Lysozyme | 12 | 11 | 11 | 7 | 2 | 0 | 9 | 0 | 0 | 2 | 1 | 4 |
| **OG0000388** | Globin Domain-Containing Protein | 27 | 10 | 7 | 9 | 0 | 0 | 2 | 0 | 0 | 0 | 0 | 0 |
| **OG0000429** | Chitin-Binding Type-4 Domain-Containing Protein | 5 | 6 | 5 | 6 | 1 | 0 | 2 | 9 | 7 | 8 | 2 | 0 |
| **OG0000161** | Chitin Binding Peritrophin-A | 31 | 21 | 12 | 19 | 0 | 0 | 0 | 1 | 0 | 0 | 0 | 0 |
| **OG0000035** | Chitinase、Chitotriosidase-1 | 24 | 31 | 23 | 18 | 7 | 3 | 4 | 12 | 15 | 15 | 6 | 0 |
| **OG0000018** | Lamin-G Domain Protein，Mucin，Chiinese | 55 | 58 | 55 | 15 | 0 | 1 | 4 | 0 | 11 | 8 | 5 | 3 |
| **OG0000311** | Glycoprotein-N-Acetylgalactosamine 3-Beta-Galactosyltransferase 1 | 9 | 13 | 10 | 9 | 4 | 2 | 5 | 3 | 3 | 1 | 2 | 0 |
| **OG0000282** | C-Type Lectin Perlucin | 17 | 10 | 15 | 7 | 0 | 1 | 2 | 1 | 11 | 0 | 0 | 0 |
| **OG0000197** | C-Type Lectin Domain-Containing Receptor 2 | 12 | 14 | 12 | 13 | 9 | 3 | 5 | 1 | 0 | 0 | 3 | 4 |
| **OG0000271** | Low Density Lipoprotein Receptor-Related Protein 5-Related | 15 | 16 | 17 | 17 | 0 | 0 | 0 | 0 | 0 | 0 | 1 | 0 |

**Supplementary Table S11** Gene families were significantly expanded in the genomes of two seep-dwelling tubeworms

| | | **P. Echinospica** | **L. Luymesi** | **R. Piscesae** | **R. Pachyptila** | **C. Teleta** | **H. Robusta** | **E. Andrei** | **L. Gigantea** | **A. Californica** | **O. Bimaculoides** | **P. Australis** | **N. Geniculatus** |
|---|---|---|---|---|---|---|---|---|---|---|---|---|---|
| **OG0000601** | Ovochymase-1 | 20 | 15 | 5 | 4 | 0 | 0 | 0 | 0 | 0 | 0 | 0 | 0 |
| **OG0000135** | Inhibitor Of Apoptosis | 14 | 16 | 10 | 9 | 8 | 0 | 10 | 9 | 1 | 1 | 3 | 11 |
| **OG0000082** | Toll-Like Receptor 4 | 38 | 43 | 21 | 13 | 0 | 0 | 0 | 0 | 0 | 0 | 0 | 0 |
| **OG0000222** | Collagen Triple Helix Repeat-Containing Protein 1 | 17 | 36 | 8 | 9 | 0 | 0 | 1 | 0 | 0 | 0 | 0 | 0 |
| **OG0000232** | Hemerythrin | 17 | 19 | 9 | 5 | 0 | 18 | 2 | 0 | 0 | 0 | 0 | 0 |
| **OG0000560** | Tyrosine-Protein Kinase | 9 | 12 | 8 | 8 | 1 | 0 | 6 | 1 | 0 | 0 | 0 | 0 |
| **OG0000632** | Thiosulfate Sulfurtransferase | 6 | 4 | 5 | 4 | 6 | 0 | 0 | 3 | 4 | 2 | 2 | 6 |
| **OG0000885** | Poly (Adp-Ribose) Glycohydrolase | 5 | 6 | 4 | 4 | 1 | 1 | 7 | 1 | 1 | 1 | 2 | 2 |
| **OG0000272** | Cartilage Intermediate Layer Protein 1 | 27 | 15 | 16 | 8 | 0 | 0 | 0 | 0 | 0 | 0 | 0 | 0 |
| **OG0000072** | Protein Mono-Adp-Ribosyltransferase Parp14 | 27 | 35 | 13 | 22 | 1 | 2 | 12 | 0 | 2 | 0 | 6 | 0 |
| **OG0000389** | IRE1 | 13 | 26 | 4 | 9 | 0 | 0 | 0 | 0 | 0 | 1 | 1 | 1 |
| **OG0000423** | Chitin-Binding Proteins | 26 | 10 | 9 | 7 | 0 | 0 | 0 | 0 | 0 | 0 | 0 | 0 |
| **OG0000210** | Cyclic Gmp-Amp Synthase | 16 | 15 | 4 | 10 | 0 | 1 | 27 | 0 | 0 | 0 | 0 | 0 |

| | | | | | | | | | | | | | |
|---|---|---|---|---|---|---|---|---|---|---|---|---|---|
| **OG0000628** | Deoxynucleoside Triphosphate Triphosphohydrolase Samhd1 | 6 | 8 | 2 | 1 | 3 | 3 | 1 | 3 | 2 | 1 | 4 | 8 |
| **OG0000891** | Docking | 10 | 7 | 4 | 5 | 1 | 0 | 2 | 1 | 1 | 1 | 1 | 2 |
| **OG0003616** | Netrin Receptor Unc5c | 3 | 13 | 0 | 1 | 0 | 0 | 0 | 0 | 0 | 0 | 0 | 0 |
| **OG0000034** | Calcium-Activated Chloride Channel Regulator 3a | 38 | 39 | 18 | 12 | 11 | 0 | 0 | 3 | 2 | 0 | 17 | 21 |
| **OG0000048** | Complement Component-Related Sushi Domain-Containing | 47 | 34 | 21 | 19 | 2 | 0 | 0 | 5 | 3 | 1 | 3 | 6 |

**Supplementary Table S12** Positively selected genes (PSGs) in *R. piscesae*

| No. | Gene ID | Gene Name | Abbreviation |
|---|---|---|---|
| **1** | Rpi\|56775 | Ras-related and estrogen-regulated growth inhibitor | *RERG* |
| **2** | Rpi\|89465 | AN1-type zinc finger protein 2B | *ZFAND2B* |
| **3** | Rpi\|46281 | NADH dehydrogenase [ubiquinone] 1 alpha subcomplex subunit 7 | *NDUFA7* |
| **4** | Rpi\|47259 | alkB homolog 2, alpha-ketoglutarate-dependent dioxygenase | *ALKBH2* |
| **5** | Rpi\|45921 | Aminoacyl tRNA synthase complex-interacting multifunctional protein 1 | *AIMP1* |
| **6** | Rpi\|52765 | Rho GTPase-activating protein 6 | *ARHGAP6* |
| **7** | Rpi\|89449 | Succinate dehydrogenase assembly factor 3 | *SDHAF3* |
| **8** | Rpi\|2573 | Derlin-1 | *DERL1* |
| **9** | Rpi\|91567 | Trifunctional enzyme subunit beta, mitochondrial | *HADHB* |
